# In vitro reconstitutions suggest a general model for paradoxical activation of ARAF, BRAF, and CRAF by diverse RAF inhibitor types that does not rely on negative allostery

**DOI:** 10.64898/2025.12.15.694443

**Authors:** Emre Tkacik, Dong Man Jang, Kayla Boxer, Byung Hak Ha, Michael J. Eck

**Affiliations:** Department of Cancer Biology, Dana-Farber Cancer Institute, Boston, MA 02215, USA; Department of Biological Chemistry and Molecular Pharmacology, Harvard Medical School, Boston, MA 02115, USA; Systems, Synthetic, and Quantitative Biology PhD Program, Harvard Medical School, Boston, Massachusetts, USA

**Author notes:** Corresponding Author: Michael J. Eck.

## Abstract

RAF kinases are central regulators of the RAS/MAP kinase pathway and important targets in cancer therapy. Paradoxically, RAF inhibitors can activate wild-type RAF signaling. Negative allostery is a central feature of the prevailing model for this phenomenon, wherein inhibitors induce RAF dimers in which inhibitor binding to one protomer promotes an active but inhibitor-resistant conformation in the other protomer. Here we systematically examined paradoxical activation of ARAF, BRAF, and CRAF using biochemical assays with isolated RAF/MEK kinase domain complexes. We found that type I and type II inhibitors induce paradoxical activation of all three isoforms, and that phosphomimetic mutation of the N-terminal acidic motif of ARAF and CRAF dramatically sensitized these isoforms to activation by type II inhibitors. The inhibition phase of paradoxical activation curves for type II inhibitors was suggestive of positive cooperativity, a finding in conflict with the prevailing model which implies negative cooperativity. In contrast to the kinase domain RAF/MEK preparations, full-length autoinhibited RAF/MEK/14-3-3 complexes were refractory to activation. Mass photometry confirmed that paradoxical activators promote BRAF dimerization. These findings support a revised model that does not rely on negative allostery. Inhibitors act on the RAS-engaged "open monomer" state to induce dimerization and activation. The open monomer and active dimer are structurally distinct species with differing affinities for inhibitor and ATP, creating a concentration window in which paradoxical activation occurs.

## Introduction

RAF family kinases (ARAF, BRAF, and CRAF) are a crucial link in the RAS/MAP kinase signaling pathway, which controls fundamental aspects of cell physiology including growth, proliferation, differentiation, and survival (1–3). RAFs are the first kinase in the three-tiered MAP kinase cascade; they phosphorylate their sole substrate MEK1/2 which in turn phosphorylates and activates ERK1/2. Mutations in BRAF are a common cause of diverse cancers, and though rare, recurrent oncogenic mutations have also been identified in ARAF and CRAF (4–6). Many inhibitors targeting BRAF and other RAF family members have been developed and several are now FDA approved for BRAF-driven cancers, in particular for malignant melanoma driven by BRAF^V600E^ (4, 7).

RAF inhibitors have long been known to induce activation of the MAP kinase pathway, leading to increased phosphorylation of ERK (8). This phenomenon, known as paradoxical activation, has important clinical ramifications. RAF inhibitors vemurafenib, dabrafenib, and encorafenib are potent and mutant-selective inhibitors of BRAF^V600E^, but they paradoxically activate wild-type RAFs, resulting in drug-induced secondary skin tumors including squamous cell carcinomas and keratoacanthomas (9, 10). To blunt this effect, these drugs are administered together with a MEK inhibitor (cobimetinib, trametinib and binimetinib, respectively) to treat BRAF-mutant melanoma (11, 12).

Vemurafenib and other BRAF^V600E^-selective RAF inhibitors are classified as type I.5 kinase inhibitors, meaning that they bind in the kinase ATP-site but extend into an allosteric pocket that requires an inactive, C-helix-out conformation of the kinase (13, 14). This binding mode affords selectivity for BRAF^V600E^ because this mutant is active as a monomer, whereas RAS-activated RAFs are dimeric. Dimerization promotes the active, C-helix-in conformation of the kinase and indeed can confer resistance to type I.5 RAF inhibitors, which are also known as RAF monomer inhibitors (14). Type I and type II RAF inhibitors also bind in the ATP-site, but are active against RAF dimers. Type I compounds bind the active conformation of the kinase, while type II inhibitors bind a specific, “DFG-out” inactive conformation of the kinase (13). The DFG motif is a three-residue sequence (Asp-Phe-Gly) in the activation segment of the kinase, and type II inhibitors bind or induce an altered conformation of this segment that is not compatible with ATP-binding or catalysis. Paradoxical activation occurs with all three types of RAF inhibitor, but is most pronounced with type I.5 inhibitors such as vemurafenib and dabrafenib (14). Paradox-breaking inhibitor PLX8394 is also a type I.5 inhibitor, but is thought to avoid paradoxical activation because it binds with sufficient potency to the C-helix-out conformation to break RAF dimers (15).

Paradoxical activation has been extensively studied. Key insights into the mechanism came with the discovery that ATP-competitive RAF inhibitors can inhibit mutant BRAF^V600E^, but at the same time enhance ERK signaling via inhibitor-induced transactivation of wild-type RAF dimers (16–18). This activation was found to require inhibitor binding at the RAF ATP site and was also dependent on a RAS-on state (16–18). RAFs are activated by RAS-driven recruitment to the membrane, which promotes catalytic activation by inducing dimerization or heterodimerization of their kinase domains (19–23). Further studies of paradoxical activation showed that certain inhibitors can stabilize a conformation of the kinase domain that promotes dimerization (24). More recently, cryo EM and crystal structures of BRAF in the autoinhibited state revealed a key role for ATP in stabilizing the inactive, monomeric conformation of the kinase domain, providing an understanding of how diverse ATP-competitive inhibitors can promote activation by displacing ATP. These and other findings have been interpreted in support of the prevailing model for paradoxical activation, which posits that inhibitor binding induces RAF dimers in which inhibitor binding to one protomer in the dimer promotes an active but inhibitor-resistant conformation in the other protomer (17, 18).

This model is seductive in that it provides a simple, unified explanation for both catalytic activation by paradoxical activators and their subsequent failure to inhibit the dimers they have induced. However, a weakness of this model is that it conflates these two distinct processes, induced dimer formation on the one hand and ineffective inhibition of these dimers on the other. While it is well-established that RAF inhibitors induce dimers by binding in the ATP-site (17, 24), it is not clear that the resulting paradoxical activity arises from half-occupied dimers or that the two protomers in a dimer differ in their affinity for drug. Type I.5 inhibitors are ineffective dimer inhibitors due to their C-helix-out binding mode, but type I and type II inhibitors also induce paradoxical activation, even though they are potent dimer inhibitors. The prevailing model implies negative cooperativity of inhibitor binding; higher affinity binding to the first site in a dimer that to the second, but type II RAF inhibitors appear to inhibit RAF dimers with positive, rather than negative, cooperativity (25, 26).

To our knowledge, paradoxical activation of individual RAF isoforms has not been previously explored at a biochemical level. Here we systematically probe the effect of structurally diverse RAF inhibitors in biochemical assays with inactive-state, monomeric preparations of ARAF, BRAF and CRAF. These kinase domain-only RAF/MEK complexes consist of their respective kinase domains bound to MEK1 and they adopt the inactive conformation observed in full-length, autoinhibited RAF complexes with MEK1 and 14-3-3. We find that type I and type II inhibitors induce paradoxical activation of all three RAF family members, and that paradoxical activation of ARAF and CRAF is primed by activating mutations in their N-terminal acidic (NtA) motifs, putative sites of tyrosine phosphorylation just N-terminal to their kinase domains. Using an *in vitro* MAP kinase pathway reconstitution, we observe that small paradoxical increases in BRAF activity can be amplified to yield substantial increases in ERK activation. Full-length autoinhibited BRAF and CRAF complexes with MEK1 and 14-3-3 are not paradoxically activated, which implies that paradoxical activation requires release of the inhibitory restraints of the 14-3-3 and RAS-binding and cysteine-rich domains (RBD/CRD). Experiments with BRAF show that, as expected, the type I and type II inhibitors studied here directly promote dimerization.

## Results

### Type I and type II RAF inhibitors paradoxically activate RAF monomers

We prepared wild-type ARAF, BRAF, and CRAF kinase domains in complex with a non-phosphorylatable variant of MEK1 (S218A/S222A, MEK1^SASA^) and tested the effect of diverse type I, type I.5 and type II RAF inhibitors on their activity in a TR-FRET based kinase assay. Co-expression of the RAF kinase domains with MEK1 allows their isolation in a monomeric state in which they adopt an inactive conformation that is essentially the same as that observed in the context of the full-length autoinhibited BRAF and CRAF complexes with MEK1 and a 14-3-3 dimer (Figure 1A). We refer to these RAF/MEK heterodimers as monomers or RAF monomer complexes because they contain a single RAF kinase domain, and unless discussing RAF/MEK binding specifically or describing new RAF preparations, refer to these samples as ARAF, BRAF, CRAF, etc. going forward, in the interest of clarity. Concentration-response curves for selected inhibitors are presented in Figure 1B and a complete set of curves is provided in Supplementary Figure 1.

**Figure 1.**
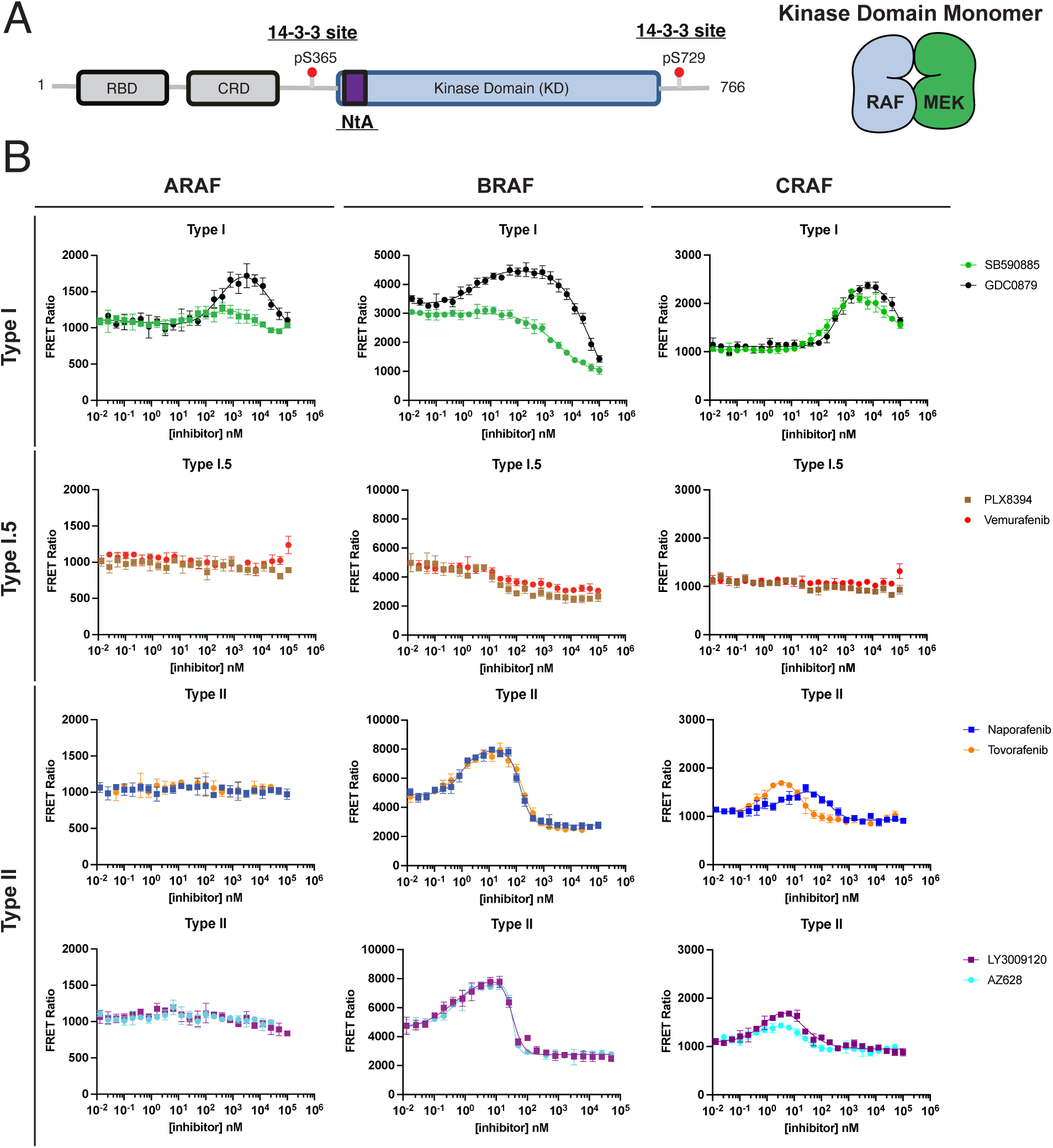
Effect of type I and type II RAF inhibitors on RAF monomer complex activity. A, Domain schematic of full-length RAF proteins, with the N-terminal acidic (NtA) motif shaded purple. Conserved 14-3-3 binding sites are indicated, with residue numbers corresponding to human BRAF. Co-expression of the RAF kinase domains with MEK1 allows for purification of the RAF/MEK1 monomer complex depicted schematically on the right and referred to as ARAF, BRAF, or CRAF in B. B, Representative concentration-response curves for the indicated type I, type I.5, and type II inhibitors with ARAF, BRAF, and CRAF monomer complexes are plotted as mean ± SD from one independent experiment performed in triplicate (n≥3).

Titration of wild-type ARAF, BRAF and CRAF monomers with type I inhibitor GDC0879 results in bell-shaped activity profiles characteristic of paradoxical activation for all three RAF isoforms (Figure 1B). Consistent with their monomeric state and our previous characterization (26, 27), these RAF monomers exhibit little (BRAF) or no (ARAF and CRAF) catalytic activity in the absence of inhibitor. ARAF and CRAF were activated to 1.65-fold and 2.5-fold above background signal, respectively, at low GDC0879 concentrations but were then inhibited at higher inhibitor concentrations (above ∼10 μM). BRAF was activated to over 150% of its basal activity at low GDC0879 concentrations and was fully inhibited at higher concentrations. We observed a similar pattern with type I inhibitor SB590885 against ARAF and CRAF, but SB590885 was less activating, elevating FRET signal to 1.2-fold above background for ARAF and 2-fold above background for CRAF (Figure 1B). Interestingly, SB590885 did not induce apparent activation in BRAF, but its IC_50_ was shifted markedly as compared with our previous measurement of its potency against 14-3-3 stabilized BRAF dimers (1.9 μM here as compared with ∼3 nM against BRAF dimers) (25). This may be a result of the superimposed effects of activation and inhibition within this concentration-response curve.

We fit the concentration-response data for these and other inhibitors with a 7-parameter bell-shaped model using GraphPad Prism to obtain activation (AC_50_) and inhibition (IC_50_) constants, as well as corresponding activation and inhibition Hill slopes (nH). These values are presented in Table 1 for the type I and type II inhibitors studied here.

**Table 1.** AC_50_, IC_50_, and Hill slopes (nH) for inhibitor titrations of ARAF, BRAF, and CRAF.*

|  |  | Activation |  | Inhibition |  |  |
| --- | --- | --- | --- | --- | --- | --- |
| Class | Inhibitor | AC<sub>50</sub> (nM) | nH | IC<sub>50</sub> (nM) | nH |  |
| Type I | GDC0879 | 286 ± 168 | 1.20 ± 0.55 | >10000 | -2.00 ± 0.71 | ARAF |
|  | SB590885 | 11.7 ± 14.3 | 0.28 ± 0.14 | >10000 | -1.11 ± 0.31 |  |
| Type I | GDC0879 | 8.40 ± 4.89 | 1.31 ± 0.46 | >10000 | -0.75 ± 0.10 | BRAF |
|  | SB590885 |  |  | 1915 ± 889 | -0.59 ± 0.19 |  |
| Type II | Tovorafenib | 0.75 ± 0.20 | 1.02 ± 0.11 | 209 ± 101 | -1.63 ± 0.10 |  |
|  | Naporaafenib | 0.76 ± 0.24 | 1.15 ± 0.23 | 148 ± 26.2 | -1.68 ± 0.35 |  |
|  | Ponatinib | 1.51 ± 0.92 | 0.97 ± 0.21 | 303 ± 131 | -1.81 ± 0.60 |  |
|  | AZ628 | 0.41 ± 0.15 | 1.30 ± 0.27 | 31.6 ± 6.74 | -2.97 ± 1.46 |  |
|  | Belvarafenib | 0.94 ± 0.66 | 1.45 ± 0.50 | 43.0 ± 3.94 | -3.49 ± 1.68 |  |
|  | LY3009120 | 0.44 ± 0.15 | 1.17 ± 0.14 | 35.7 ± 2.71 | -2.81 ± 0.94 |  |
| Type I | GDC0879 | 399 ± 96.1 | 1.49 ± 0.11 | >10000 | -1.50 ± 0.29 | CRAF |
|  | SB590885 | 242 ± 19.5 | 1.32 ± 0.24 | >10000 | -1.40 ± 0.11 |  |
| Type II | Tovorafenib | 0.53 ± 0.09 | 1.41 ± 0.49 | 26.0 ± 4.77 | -1.63 ± 0.33 |  |
|  | Naporaafenib | 3.06 ± 1.56 | 0.88 ± 0.15 | 285 ± 67.8 | -2.05 ± 0.51 |  |
|  | Ponatinib | 1.63 ± 0.55 | 2.28 ± 0.88 | 81.9 ± 19.4 | -1.33 ± 0.12 |  |
|  | AZ628 | 0.27 ± 0.10 | 1.42 ± 0.43 | 23.4 ± 4.58 | -2.16 ± 0.57 |  |
|  | Belvarafenib | 1.31 ± 0.68 | 2.21 ± 1.66 | 17.9 ± 6.82 | -1.76 ± 0.36 |  |
|  | LY3009120 | 0.32 ± 0.10 | 1.23 ± 0.37 | 25.0 ± 3.62 | -2.14 ± 0.09 |  |

\*Data are reported as mean ± SD of at least three independent experiments, each of which was performed in triplicate (n≥3).

Type I.5 inhibitors did not induce activation in this biochemical assay (Figure 1B, Supplementary Figure 1). This is not unexpected, considering that they require a C-helix-out binding mode that disfavors dimerization (13, 28). Type 1.5 compounds vemurafenib and dabrafenib are known to induce paradoxical activation in cells, where the effects of RAS and membrane engagement together with 14-3-3 binding can act in concert with inhibitor binding to promote dimerization and activation (14, 17, 18, 29). These features are absent in our biochemical assay and thus we do not observe RAF catalytic activation or dimerization (see below) in the presence of these compounds.

With type II inhibitors, we observed a unique behavior. All six type II inhibitors tested induced marked activation of BRAF and CRAF, but not ARAF (Figure 1B, Supplementary Figure 1). Activation of BRAF was particularly pronounced, with maximal activation of approximately 200% of basal activity. For each of the six inhibitors tested, the activation phase was followed by a steep inhibition phase. Hill constants for the inhibition phase obtained with the curve fitting approach outlined above ranged from -1.6 to -3.5 for naporafenib and belvarafenib, respectively, reflective of strong positive cooperativity as we have previously observed for inhibition of 14-3-3 bound RAF dimers by type II inhibitors (25, 26). Activation of CRAF was somewhat less pronounced and inhibition was less steep, with Hill constants indicative of little to no positive cooperativity (Figure 1B, Table 1).

#### Inhibitor-induced paradoxical activation is amplified in pathway reconstitutions

We next examined paradoxical activation by selected inhibitors in a biochemical reconstitution of the MAP kinase pathway. This cascade assay contains the BRAF monomer complex described above, along with wild-type MEK1 and ERK2 and uses TR-FRET to measure the increase in ERK2 phosphorylation. MEK phosphorylation was measured in parallel reactions with the TR-FRET assay employed in Figure 1. As in the assay above, type I inhibitor GDC0879 and type II inhibitors naporafenib and AZ628 induced paradoxical activation, while type I.5 inhibitors vemurafenib and PLX8394 did not (Figure 2A,B). The concentration ranges over which each inhibitor induced activation were essentially the same both in both pMEK and pERK assay systems and, as might be expected in a cascade assay where signaling is amplified down the pathway, the magnitude of the increase in ERK phosphorylation was greater than that observed for MEK. For example, naporafenib induced a ∼1.5 fold increase in pMEK, but more than a 3-fold increase in pERK.

**Figure 2.**
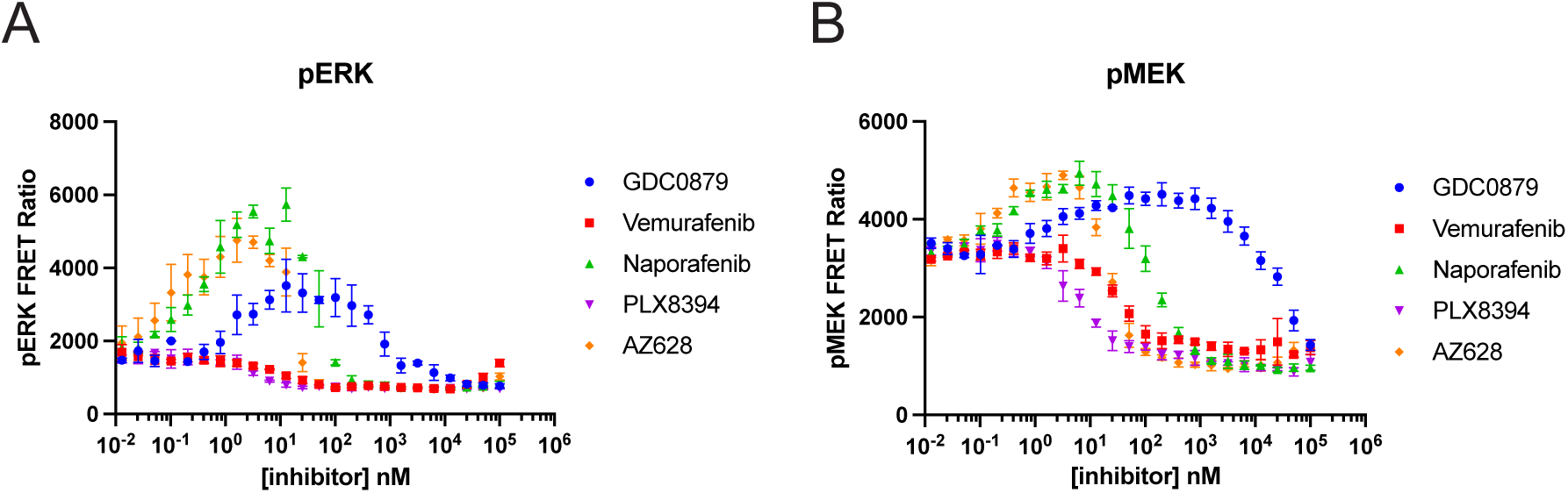
Effect of type I and type II inhibitors in an *in vitro* MAP kinase pathway reconstitution with BRAF, MEK1 and ERK2. A, Paradoxical activation of BRAF as measured by changes in ERK phosphorylation. B, Paradoxical activation of BRAF as measured by changes in MEK phosphorylation. Phospho-MEK and Phospho-ERK were measured in parallel reactions. Data are plotted as mean ± SD from one of three independent experiments performed in triplicate (n=3).

#### Full-length autoinhibited RAF/MEK/14-3-3 complexes are not paradoxically activated

We also tested the ability of selected inhibitors to paradoxically activate full-length autoinhibited BRAF/MEK1/14-3-3 and CRAF/MEK1/14-3-3 complexes (30, 31). In the intact autoinhibited complexes, the conformation of the RAF/MEK kinase domain region is essentially the same as in the isolated monomer complexes above, but the RAF kinase domains are protected from dimerization by interactions with the 14-3-3 dimer and the N-terminal regulatory domains of the kinase (the RAS-binding and cysteine-rich domains, RBD/CRD, Supplemental Figure 2A). Unlike the isolated kinase domain complexes, the full-length BRAF and CRAF complexes were not activated by any of the inhibitors tested (Supplemental Figure 2B,C). Thus while type I and type II inhibitors paradoxically activate isolated RAF monomer complexes, which lack the protective interactions with the RBD/CRD and 14-3-3 dimer, they do not activate the fully autoinhibited RAF/MEK/14-3-3 complexes. This biochemical finding is consistent with studies of paradoxical activation in a cellular context, where a RAS-on state is required for the effect. RAS-driven membrane recruitment of RAFs is expected to induce “opening” of the autoinhibited complex, removing the steric restraints of the CRD and 14-3-3 dimer to allow dimerization.

#### Mutation of the NtA motif primes ARAF and CRAF for paradoxical activation

RAFs contain a sequence at the N-terminus of the kinase domain known as the N-terminal acidic (NtA) motif. In BRAF, this motif has the sequence “SSDD” but in ARAF and CRAF the aspartic acid residues are replaced by tyrosine (with NtA sequences SGYY and SSYY in ARAF and CRAF, respectively). ARAF and CRAF are thought to require phosphorylation of the second tyrosine in this motif for full activity, and mutation of this region to match the SSDD sequence of BRAF has been previously shown to increase their activity, presumably by obviating the need for tyrosine phosphorylation of their native NtA motifs (32–34). For this reason, we also prepared and tested ARAF and CRAF monomer complexes with their respective NtA motifs mutated to SSDD similarly to wild-type ARAF and CRAF (see Materials and Methods). The basal activity of both ARAF^SSDD^ and CRAF^SSDD^ was modestly increased as compared with their wild-type counterparts (Figure 3A).

**Figure 3.**
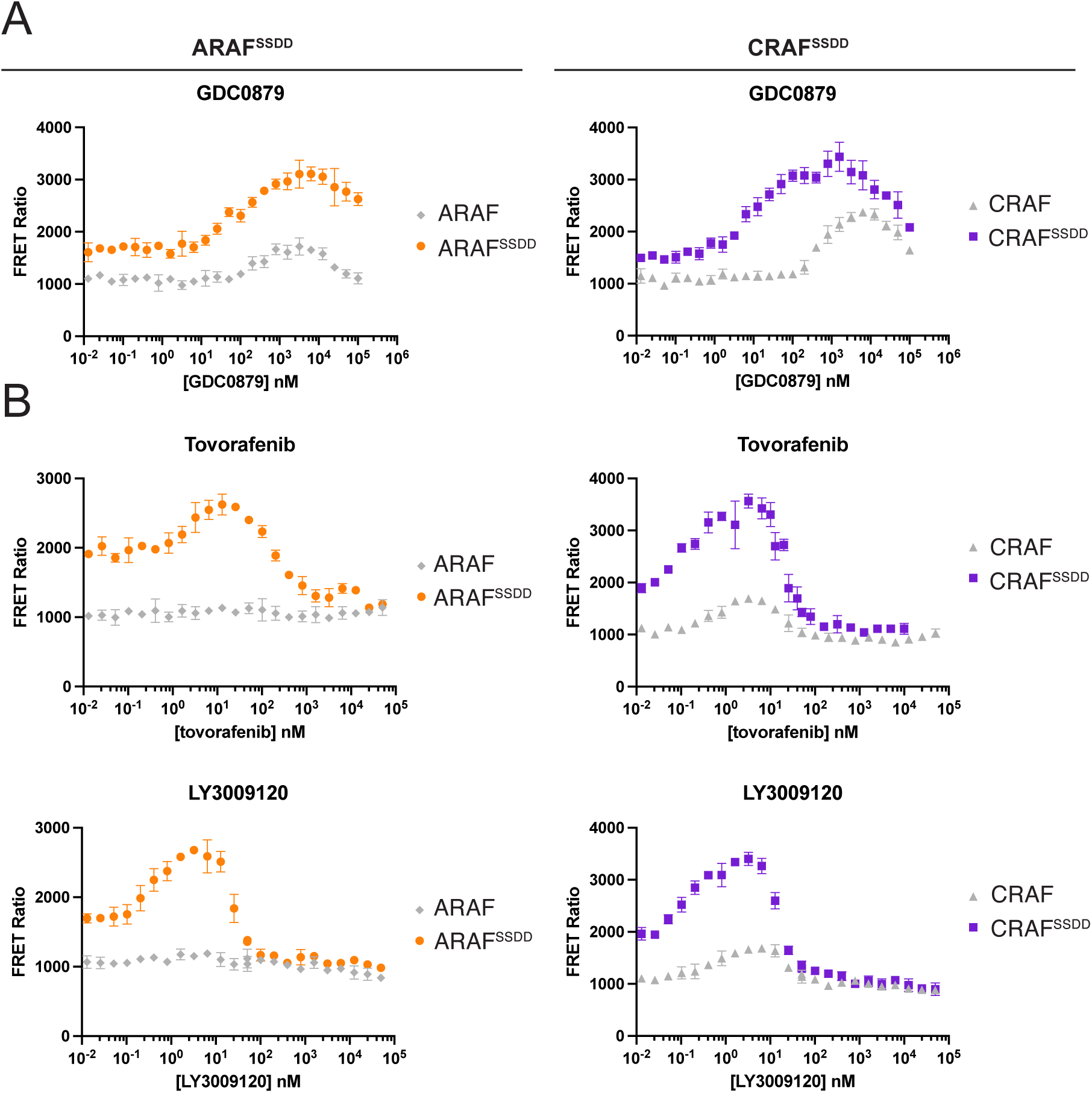
Phosphomimetic mutations of the NtA motif prime ARAF and CRAF monomer complexes for paradoxical activation. A, Concentration-response curves for titration of ARAF^SSDD^/MEK1 (ARAF^SSDD^) and CRAF^SSDD^/MEK1 (CRAF^SSDD^) with type I inhibitor GDC0879. B, Concentration-response curves for titration of ARAF^SSDD^ and CRAF^SSDD^ with type II inhibitors tovorafenib and LY3009120. Data for the wild-type ARAF and CRAF are reproduced from Figure 1 to facilitate comparison and are plotted in gray. Curves for additional type I and type II inhibitors are shown in Supplemental Figure 3. Data are plotted as mean ± SD from one independent experiment performed in triplicate.

The paradoxical activation effect for both type I and type II inhibitor titrations was more striking on these SSDD variants, in particular for type II inhibitors (Figure 3A,B). While wild type ARAF was not activated by type II inhibitors, ARAF^SSDD^ exhibited prominent bell-shaped concentration-response curves similar to those observed with BRAF (Figure 3B, Table 2, Supplemental Figure 3A). For both type I and type II inhibitors, activation of CRAF^SSDD^ was much more prominent than for wild type CRAF and was evident at lower inhibitor concentrations (Figure 3A,B, Supplemental Figure 3B). For type II inhibitors in general, Hill slopes for the inhibition portion of the curves were steeper, indicative of positive cooperativity as we have observed for inhibition of 14-3-3-bound CRAF dimers by these compounds (Figure 3B, Table 2, Supplemental Figure 3B).

**Table 2.** AC_50_, IC_50_, and Hill slopes (nH) for inhibitor titrations of ARAF^SSDD^ and CRAF^SSDD^.*

|  |  | Activation |  | Inhibition |  |  |
| --- | --- | --- | --- | --- | --- | --- |
| Class | Inhibitor | AC<sub>50</sub> (nM) | nH | IC<sub>50</sub> (nM) | nH |  |
| Type I | GDC0879 | 129 ± 90.3 | 0.79 ± 0.24 | >10000 | -1.84 ± 0.93 | ARAF<sup>SSDD</sup> |
|  | SB590885 | 7.62 ± 2.83 | 1.16 ± 0.03 | >10000 | -1.11 ± 0.63 |  |
| Type II | Tovorafenib | 1.33 ± 0.35 | 1.40 ± 0.55 | 154 ± 147 | -1.85 ± 1.53 |  |
|  | Naporafenib | 13.3 ± 5.15 | 1.71 ± 1.04 | 673 ± 488 | -1.26 ± 0.58 |  |
|  | Ponatinib | 52.2 ± 9.18 | 0.87 ± 0.50 | 517 ± 99.8 | -1.13 ± 0.39 |  |
|  | AZ628 | 1.72 ± 1.24 | 1.23 ± 0.53 | 44.4 ± 17.6 | -1.43 ± 0.12 |  |
|  | Belvarafenib | 1.59 ± 0.50 | 1.44 ± 0.48 | 94.6 ± 6.70 | -1.79 ± 0.07 |  |
|  | LY3009120 | 0.48 ± 0.13 | 1.38 ± 0.26 | 28.3 ± 3.73 | -2.21 ± 0.44 |  |
| Type I | GDC0879 | 20.1 ± 9.53 | 0.84 ± 0.17 | >10000 | -1.11 ± 0.33 | CRAF<sup>SSDD</sup> |
|  | SB590885 | 3.46 ± 1.49 | 1.06 ± 0.22 | >10000 | -0.87 ± 0.06 |  |
| Type II | Tovorafenib | 0.13 ± 0.10 | 0.70 ± 0.32 | 23.2 ± 7.93 | -1.81 ± 0.17 |  |
|  | Naporafenib | 0.06 ± 0.04 | 0.84 ± 0.79 | 79.2 ± 24.1 | -1.48 ± 0.65 |  |
|  | Ponatinib | 0.08 ± 0.07 | 0.75 ± 0.78 | 70.4 ± 25.5 | -1.93 ± 0.68 |  |
|  | AZ628 | 0.02 ± 0.03 | 0.60 ± 0.39 | 27.5 ± 4.76 | -3.25 ± 0.77 |  |
|  | Belvarafenib | 0.07 ± 0.06 | 0.46 ± 0.13 | 34.6 ± 5.15 | -2.39 ± 0.36 |  |
|  | LY3009120 | 0.11 ± 0.10 | 0.67 ± 0.17 | 26.3 ± 12.7 | -2.26 ± 0.30 |  |

\*Data are reported as mean ± standard deviation of at least three independent experiments, each of which was performed in triplicate (n≥3), except for the belvarafenib experiment with CRAF<sup>SSDD</sup>, which was performed twice in triplicate (n=2).

#### ATP concentration modulates activation by type I inhibitors

Considering the central role of ATP in maintaining RAF autoinhibition and its displacement in paradoxical activation (35), we were interested in directly assessing the effect of ATP concentration on paradoxical activation. We titrated the BRAF monomer complex with type I and type II inhibitors at ATP concentrations of 10 µM, 100 µM, and 1000 µM and measured the resulting MEK phosphorylation. With type I inhibitors GDC0879 and SB590885, the apparent magnitude of the paradoxical activation effect was increased at higher ATP concentrations and peak activity was shifted to higher inhibitor concentrations, as is most clearly seen in the normalized concentration-response curves (Figure 4A,B). We attribute the more marked activation at higher ATP concentrations to tighter autoinhibition (lower activity) of the monomer complex under these conditions. In addition, the inhibition phases of the concentration-response curves were markedly right shifted, as expected for ATP-competitive inhibition (Figure 4A,B, Supplemental Table 1). The shift in IC_50_ values was dramatic, roughly 10-fold for each 10-fold increase in ATP concentration. We note that this is as expected from the Cheng-Prusoff relation (36).

**Figure 4.**
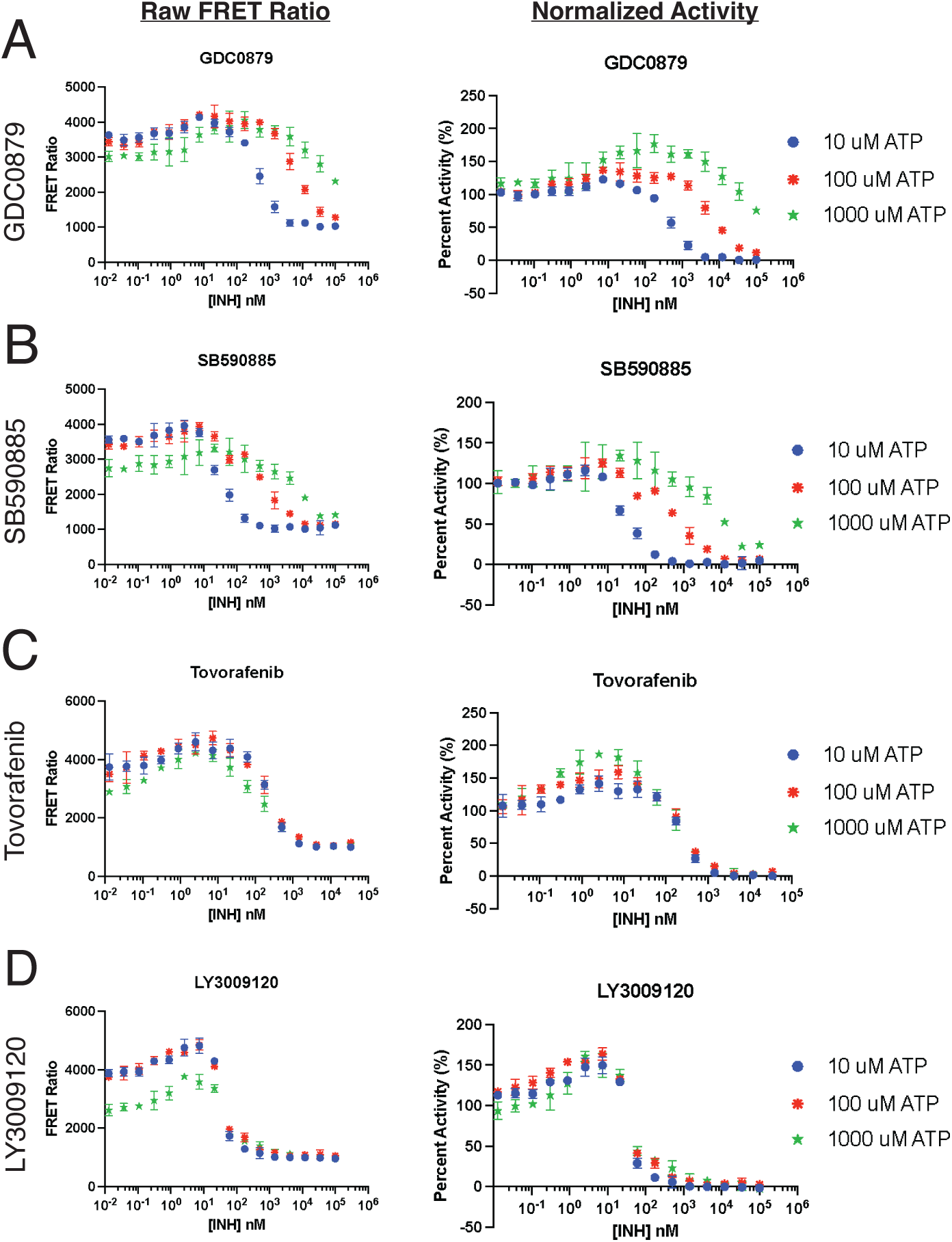
Effect of ATP concentration on paradoxical activation of BRAF by type I and type II inhibitors. A-D, Concentration-response curves measuring MEK phosphorylation upon treatment of the BRAF monomer complex with increasing concentrations of the indicated inhibitor at ATP concentrations of 10 μM (blue), 100 μM (red), and 1000 μM (green). MEK phosphorylation activity is plotted as the raw FRET ratio in the panels on the left, and is replotted on the right as percent activity, normalized to the activity in the absence of inhibitor at each ATP concentration. Data are plotted as mean ± SD from one independent experiment performed in triplicate (n=3).

With type II inhibitors, ATP concentration had surprisingly little effect on the observed concentration-response profiles (Figure 4C,D). For tovorafenib and LY3009120, concentrations at which peak activity was observed were essentially unchanged, as were AC_50_ and IC_50_ values (Supplemental Table 1). These compounds are clearly ATP-competitive in that they bind in the ATP-site, but they induce a DFG-out conformation of the kinase that is not compatible with ATP-binding, giving rise to this apparent non-competitive behavior. While this is not a universal feature of type II kinase inhibition, it was evident it in our recent studies of inhibition of 14-3-3-bound RAF dimers with these compounds (25). We obtained comparable results with type II inhibitors ponatinib, AZ628, and belvarafenib (Supplemental Figure 4, Supplemental Table 1).

#### Measuring BRAF dimerization with single molecule mass photometry

We employed mass photometry (MP) to directly measure BRAF dimerization and association with MEK in the presence and absence of inhibitors. MP detects individual macromolecules or complexes as they bind and unbind to a glass coverslip mounted atop an inverted video microscope. Binding events to the cover slip result in changes in light scattering that are directly proportional to the size of the incident macromolecule. MP can thus reveal the proportions of various BRAF:MEK complexes present in the solution examined. In addition to our BRAF monomer complex (the BRAF:MEK kinase heterodimer, 80 kDa) and individual BRAF and MEK monomers (∼35 kDa and ∼45 kDa, respectively), we can also expect to observe MEK:BRAF:BRAF:MEK heterotetramers (∼160 kDa) and MEK:BRAF:BRAF complexes lacking one MEK (∼115 kDa), both of which we have been observed in structural studies of BRAF/MEK complexes (30). Owing to their similar size, MP cannot discriminate between BRAF:MEK and BRAF:BRAF dimers, nor between BRAF and MEK monomers. Our analysis takes these ambiguities into account, as described in detail in *Materials and Methods*.

A histogram showing the distribution of masses present in a BRAF:MEK sample in the absence of inhibitor at 32 nM BRAF with 10 μM ATPγS is shown in Figure 5A. Under these conditions, most counts fall within the 45 kDa and 80 kDa mass distribution ranges, while smaller proportions lie in the ∼115 kDa and ∼160 kDa ranges representing BRAF dimers. Given the masses of the component species, this distribution can be fit to obtain the relative abundance of each (Figure 5A). This information, together with knowledge of total protein concentration, allows calculation of K_d_ values for the various species as illustrated in the scheme in Figure 5B. Despite the limitations of this approach as noted above, the K_d_ values obtained are consistent with prior studies of RAF homodimerization and RAF:MEK heterodimerization (24, 27, 28, 37).

**Figure 5.**
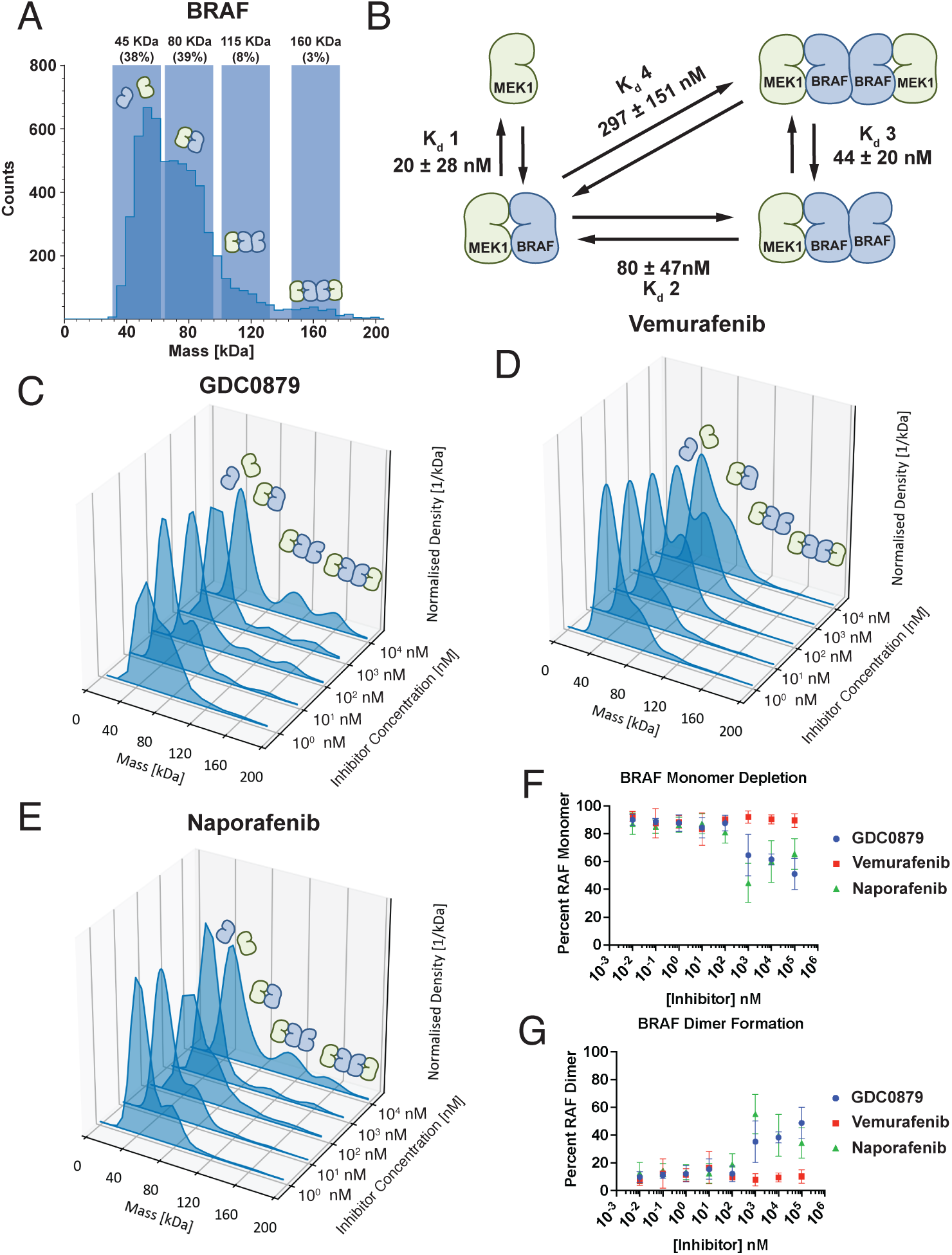
Mass photometry analysis of the oligomeric state of BRAF and MEK1. *A*, MP mass distributions of the BRAF/MEK1 complex (32 nM) in the presence 10 μM ATPγS and without addition of inhibitor. *B*, “Single-shot” binding affinities of BRAF and MEK1 interactions in the absence of inhibitor, calculated as mean ± SD from five independent MP experiments (n = 5). A representative experiment is shown in panel A. *C*, Mass distributions of the BRAF/MEK1 complex (32 nM) in ATPγS (10 μM) at increasing concentrations of inhibitor GDC0879. *D*, Mass distributions of the BRAF/MEK1 complex (32 nM) in ATPγS (10 μM) at increasing concentrations of inhibitor vemurafenib. *E*, Mass distributions of the BRAF/MEK1 complex (32 nM) in ATPγS (10 μM) at increasing concentrations of inhibitor naporafenib. For experiments in C, D, and E, inhibitor concentrations from 10^-2^ nM to 10^5^ nM were tested in three independent experiments (n=3) per inhibitor concentration, but only representative experiments from 10^0^ nM to 10^4^ nM are illustrated here. See also Supplemental Tables 2 and 3. *F*, BRAF monomer depletion plotted across inhibitor concentrations, as determined by relative distributions of counts from C, D, and E. *G*, BRAF dimer formation plotted across inhibitor concentrations, as determined by relative distributions of counts from C, D, and E. In F and G, the 45 kDa and 80 kDa peaks are summed to obtain the percent BRAF monomer, and the 115 kDa and 160 kDa peaks are summed to obtain percent BRAF dimer.

**Table 3.** MP analysis of BRAF (R) and MEK1 (M) interactions at 10 μM inhibitor.*

|  | **K<sub>d</sub> (nM)** |  |  |  |
| --- | --- | --- | --- | --- |
|  | **R-M** | **RM-R** | **MRR-M** | **RM-RM** |
| **GDC0879** | 55 ± 15 | 21 ± 6 | 22 ± 3 | 9 ± 5 |
| **Vemurafenib** | 13 ± 8 | 169 ± 76 | 179 ± 18 | >2000 |
| **Naporaafenib** | 61 ± 38 | 19 ± 11 | 58 ± 37 | 20 ± 14 |
| **No inhibitor** | 20 ± 29 | 79 ± 47 | 44 ± 20 | 297 ± 151 |

\*Interactions correspond to the scheme shown in Fig. 5B. K<sub>d</sub> values are calculated as mean ± SD from at least three independent experiment (n ≥ 3). Values obtained in the absence of inhibitor from Fig. 5B are provided for reference.

Having established the utility of this approach, we applied it to measure the effect of inhibitors on the oligomeric state of BRAF and MEK1. We recorded MP data at a broad range of concentrations of a type I (GDC0879, Figure 5C), type I.5 (vemurafenib, Figure 5D) and type II inhibitor (naporafenib, Figure 5E). The BRAF:MEK1 complex (32 nM) was maintained in the presence of ATP analog ATPγS (10 μM) in these inhibitor titrations. As with the inhibitor-free data, we fit the resulting mass distributions to obtain the concentrations of each BRAF:MEK species and in turn the relevant dissociation constants in the presence of each inhibitor. The resulting K_d_ values for the inhibitor-free and 10 μM inhibitor concentrations (where shifts in K_d_ generally plateaued) are shown in Table 3, and complete data across all inhibitor concentrations is provided in Supplemental Table 2. As can be appreciated from Table 3, both GDC0879 and naporafenib markedly decreased dissociation constants for BRAF dimers (reflecting higher affinity dimerization), irrespective of MEK association, and had little effect on dissociation constants for MEK. The relative fractions of RAF monomers and dimers as a function of inhibitor concentration is plotted in Figure 5F and G, respectively, and are also provided in Supplementary Table 3. For GDC0879 and naporafenib, the fraction of BRAF dimers (including both the 115 kDa and 160 kDa species) increases markedly at higher inhibitor concentrations, accompanied by a corresponding decrease in the fraction of BRAF monomers (the 80 kDa species). By contrast, vemurafenib had little effect on the monomer/dimer equilibrium and the resulting dissociation constants reflected a modest increase in K_d_ for BRAF dimerization, albeit with large errors due the vanishingly low concentration of dimers observed in the presence of this type I.5 inhibitor (Figure 5F,G and Table 3).

## Discussion

In this study, we show that paradoxical activation can be recapitulated *in vitro* for all three RAF isoforms for both type I and type II RAF inhibitors. In systematically examining paradoxical activation with isolated RAF kinase domain monomers, we found that type I and type II inhibitors induce paradoxical activation of all three RAF isoforms, though ARAF requires priming by of the NtA motif for activation by type II inhibitors and the magnitude and inhibitor-class effects differed among isoforms. Wild-type BRAF was activated by both type I and type II inhibitors, with type II compounds producing particularly pronounced effects. Wild-type CRAF showed moderate activation by both inhibitor classes, while ARAF was activated only by type I inhibitors. Mutation of the NtA motif of ARAF and CRAF to the “phosphomimetic” SSDD sequence dramatically sensitized them to paradoxical activation by type II inhibitors, implicating this regulatory element as a key determinant of this effect. Type I.5 inhibitors produced no activation in these biochemical assays, consistent with their C-helix-out binding mode that disfavors dimerization. Full-length autoinhibited RAF/MEK/14-3-3 complexes were not susceptible to paradoxical activation, demonstrating that release from autoinhibitory interactions is required for the phenomenon. Mass photometry experiments confirmed that type I and type II, but not type I.5, inhibitors promote BRAF dimerization, and pathway reconstitution assays revealed that modest increases in RAF activity can be amplified downstream to yield substantial increases in ERK phosphorylation.

Early studies of ERK-pathway activation by RAF inhibitors established inhibitor-induced RAF dimerization as a fundamental mechanism underlying paradoxical activation (17, 18), with the effect driven by a conformational effect of the inhibitor on the RAF kinase domain (24).

Taken in isolation, this mechanism appears to pose a conundrum – why is an inhibitor-induced dimer active? The second component of the prevailing model for paradoxical activation – that inhibitor binding to one protomer of the dimer promotes an active but inhibitor resistant conformation in the other – provides an appealing resolution to the apparent puzzle. This negative-cooperativity or “negative allostery” model has its roots in an elegant chemical-genetic experiment. By introducing a cysteine mutation in CRAF which rendered it sensitive to an irreversible inhibitor and co-expressing it with wild type CRAF or BRAF, Poulikakos et al. showed that the covalent inhibitor (JAB34, a covalent EGFR inhibitor that does not inhibit WT

RAF) could induce dimerization and transactivation of the wild-type dimerization partner by binding to the sensitized (cysteine-containing) subunit of a CRAF homodimer or CRAF/BRAF heterodimer (17). While this experiment cleanly demonstrates inhibitor-induced transactivation of RAF dimers, it is important to recognize that the differential inhibitor sensitivity of the two subunits in this experiment is artificial – it is engineered rather than induced by inhibitor binding as the negative allostery model proposes.

Our results suggest that negative allostery (or negative cooperativity) is not a requisite feature of paradoxical activation. The type I and type II inhibitors studied here induce RAF dimers and exhibit paradoxical activation but do so without evidence of negative cooperativity, nor do they appear to inhibit intentionally engineered RAF dimers with negative cooperativity (25). Indeed, type II inhibitors exhibit apparent positive cooperativity while type I inhibitors are non-cooperative inhibitors of RAF dimers (25). On the other hand, type I.5 inhibitors may inhibit RAF with negative cooperativity, although in our view this is far from clear. Evidence cited in support of the negative allostery model includes crystal structures that show vemurafenib or other type I.5 RAF inhibitors bound to only one protomer of a RAF dimer with an inactive C-helix-out conformation, with the opposite protomer adopting an active, inhibitor-free conformation (38). In inhibitor studies with 14-3-3-stabilized RAF dimers, concentration response curves for selected type I.5 inhibitors including vemurafenib are well-fit with Hill coefficients consistent with negative cooperativity (25). Finally, simple thermodynamic modeling confirms that negative allostery can give rise to paradoxical activation, with paradoxical activity arising from half-occupied RAF dimers, so long as inhibitor binding promotes RAF dimerization (39–41).

We find this evidence suggestive, but not conclusive. Other RAF crystal structures with type I.5 inhibitors reveal symmetric dimers, with inhibitor occupying both active sites (PDB ID: 5csw) (42). Also, differences in inhibitor occupancy can arise due to interactions in the crystal lattice. Considering that the conformation required for binding of type I.5 inhibitors destabilizes RAF dimers, it is unclear how an inhibitor binding to one protomer would be able to transmit an allosteric change to the opposite protomer, if that inhibitor’s binding causes the existing dimer to dissociate. Furthermore, the complex effects of type I.5 inhibitors on dimer stability and the clear resistance of active RAF dimers to these inhibitors complicates interpretation of inhibition data – weak or incomplete inhibition of an enzyme can be difficult to discern from true negative cooperativity (43). As we discuss below, the clear resistance of RAF dimers to type I.5 inhibitors is alone sufficient to explain their ineffective inhibition during paradoxical activation, without invoking negative allostery.

For the most part, our biochemical findings parallel observations in previous cellular studies. Type I and type II inhibitors are well known to induce paradoxical ERK pathway activation (17, 18, 24, 39, 44), and ARAF and CRAF as well as BRAF have been shown to be direct targets for paradoxical activation in cellular contexts (17, 45, 46). The phenomenon has been shown to require a RAS-on state (16–18), which promotes recruitment of RAF to the plasma membrane (2, 23, 47). Membrane recruitment is thought to promote release of autoinhibitory interactions, leading to an “open monomer” state that is an intermediate in the activation pathway (29). Consistent with cellular studies, we find that the isolated RAF/MEK kinase monomer complex, which is a proxy for the open monomer, is susceptible to paradoxical activation whereas the full-length autoinhibited RAF/MEK/14-3-3 complex is not. RAS-driven membrane recruitment is thought to be required for phosphorylation of the NtA motif (48–50), and there is evidence that the NtA motif is phosphorylated in an inhibitor-dependent fashion (18). Interestingly, we find that activating mutations in this motif enhance paradoxical activation of ARAF and CRAF. In prior studies, we found that pre-formed 14-3-3- stabilized RAF dimers are not further activated by inhibitors (25), further pointing to the open monomer as the relevant target in paradoxical activation. One obvious disconnect in our findings as compared with cellular studies is the lack of paradoxical activation by type I.5 inhibitors in our biochemical system. As we noted above, this is likely due to the lack of 14-3-3 binding and membrane association in our simplified biochemical system.

Our findings here, in light of structural studies of RAF complexes and prior cellular investigations of paradoxical activation, lead us to a model for paradoxical activation that does not rely on negative allostery and is consistent with activation by diverse inhibitor classes. In this model, the open monomer complex is the target of inhibitor-induced paradoxical activation (Figure 6). Binding of ATP to the RAF active site stabilizes the inactive conformation of the open monomer, which disfavors dimerization. Displacement of ATP by an ATP-competitive inhibitor, irrespective of class, alters the relative N- and C-lobe orientations of the kinase to promote dimerization (30, 35). Once dimerized, inhibitor dissociation from one or both sides of the dimer would allow phosphorylation and activation of MEK.

**Figure 6.**
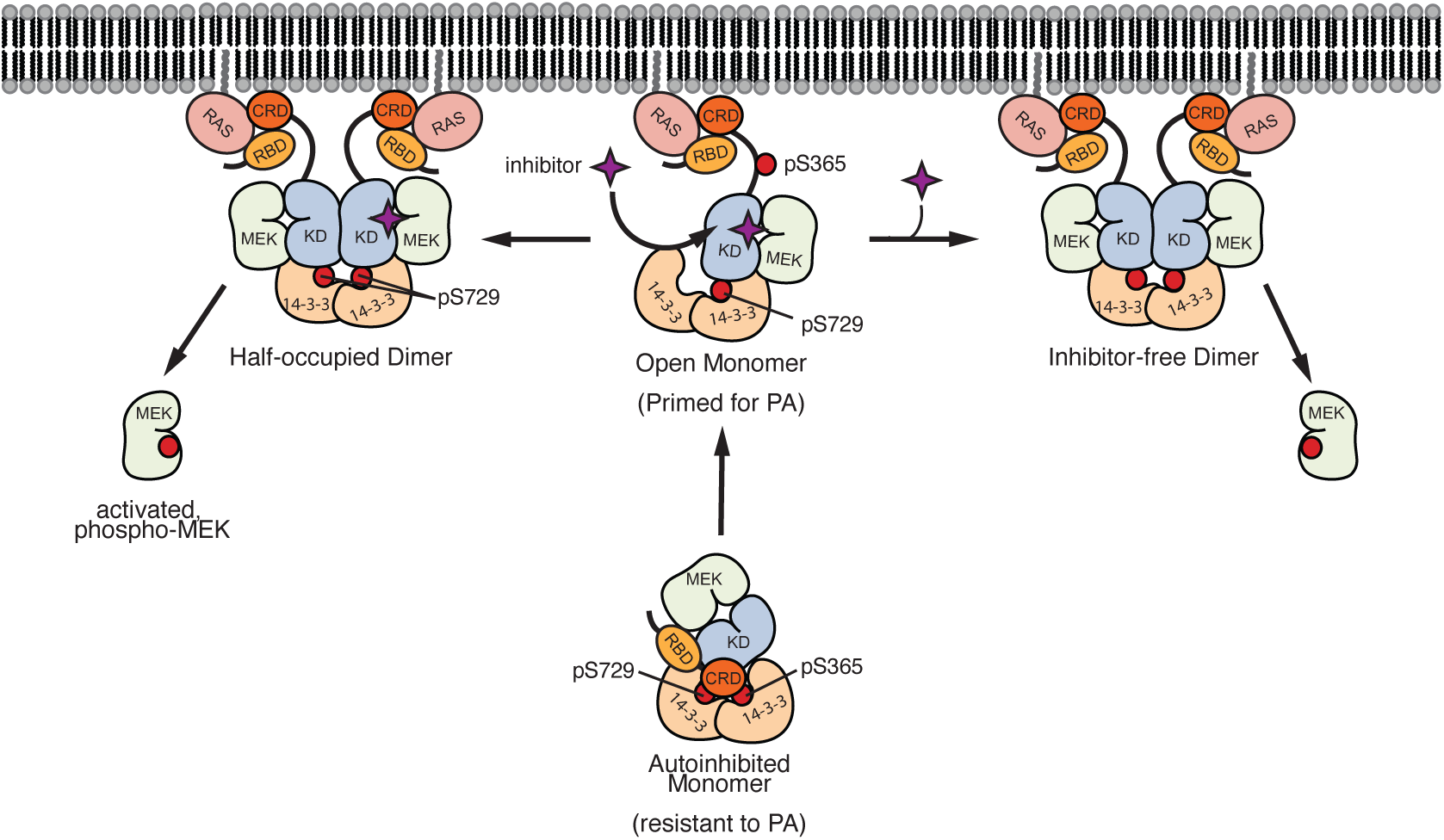
A general model for inhibitor-induced paradoxical activation of RAF signaling. In the resting state, autoinhibitory interactions between the RAS binding domain (RBD), cysteine-rich domain (CRD), and RAF kinase domain (KD) maintain RAF/MEK/14-3-3 complexes as a monomer in the cytosol. We find that this species is resistant to inhibitor-induced activation. RAS-driven recruitment to the membrane promotes opening of this complex, forming an “open” monomer that is not yet dimerized and active, but, unlike the autoinhibited monomer is susceptible to paradoxical activation (PA). Inhibitor binding to this open monomer causes paradoxical activation by displacing ATP to promote RAF dimerization, leading to the formation of signaling competent dimers. Dissociation of inhibitor from one or both sides of the dimer, or formation of a dimer with an inhibitor-free RAF, allows phosphorylation and activation of MEK. In this model, the open monomer is the target for activation, while the 14-3-3-bound dimer is the target for inhibition. Differing affinities of these distinct species for drug and for ATP can be expected to leave a window in which paradoxical activation is observed for any inhibitor that is more potent at displacing ATP from the open monomer than from the active dimer.

How does this model address the “conundrum” of how an inhibitor-induced dimer can nevertheless be active? Inhibitor-induced activation and inhibition act on distinct species – activation on the open monomer and inhibition on the 14-3-3-stabilized dimer. Each of these have their own affinities for ATP and competing drug. Provided that an inhibitor is more potent at displacing ATP from the open monomer than from the active dimer, there will be a concentration window in which paradoxical activation can be observed. It is important to note that this model presumes a lack of microscopic reversibility in this system, otherwise it would violate thermodynamic detailed balance relationships (41). Dimerization involves not only a change in oligomeric state but also a change in subunit stoichiometry – one 14-3-3 dimer per RAF dimer versus one 14-3-3 dimer per RAF monomer in the open monomer state – and it also involves changes in phosphorylation state, including dephosphorylation of the inhibitory S365 site (S259 in CRAF) by the SHOC2 phosphatase complex (51, 52). Either or both of these factors could be expected to impede rapid reversibility of dimerization. We recognize that these cellular effects are not at play in the simplified biochemical systems we employ here, where we nevertheless observe paradoxical activation across inhibitor chemotypes. In this regard, we note that we are studying inhibitor-induced paradoxical activation in end-point enzyme inhibition assays, and not under equilibrium conditions where microscopic reversibility applies.

To summarize, our findings reveal that each RAF isoform is differentially susceptible to paradoxical activation, both in the intensity of activation observed, and in the inhibitors that are able to induce this activation, with BRAF being both the most susceptible in magnitude and number of inhibitors. We find that mutation of the ARAF or CRAF NtA motif to match the SSDD sequence in BRAF increases the sensitivity of these isoforms to paradoxical activation, cementing this regulatory sequence as a key modulator of this effect. For type II inhibitors, the steep slope of the inhibition phase of the paradoxical activation curves indicates positive cooperativity, while type I inhibitors appear to be non-cooperative. Using mass photometry, we confirmed that inhibitors that exhibit paradoxical activation of BRAF *in vitro* also induce BRAF dimerization *in vitro*. Finally, we see that while RAF monomers are susceptible to paradoxical activation, RAF/MEK/14-3-3 autoinhibited complexes are not paradoxically activated under analogous conditions. Considering these findings and prior studies of the phenomenon, we outline a general model for paradoxical activation that applies across inhibitor chemotypes and does not rely on negative allostery.

## Materials and Methods

### Protein expression and purification

All the proteins used in both the TR-FRET kinase assay as well as for collecting single molecule FRET measurements were expressed and purified as described in previously (25). Briefly, truncated RAF kinase domain constructs, lacking 14-3-3 binding motifs, were co-expressed and purified in insect cells, eluting from gel filtration as RAF/MEK heterodimers with low levels of innate kinase activity. We refer to these RAF/MEK heterodimers as RAF monomers because RAF is in its inactive monomeric state. For full details, please refer to Tkacik et al (25).

BRAF monomers were obtained by co-expressing his_6_-BRAF^445-723^-intein (BRAF) and full length MEK1^S218A/S222A^ (MEK1^SASA^) constructs in SF9 insect cells, purifiying by Ni-NTA, then incubating with 50 mM MESNA overnight. Cleaved protein was applied to Chitin beads, and the flowthrough was concentrated to 1 ml and injected onto a Superdex 200 10/300 (GE Healthcare Life Sciences) sizing column. ARAF and CRAF monomers were purified by co-expressing the MEK1^SASA^ construct with his_6_-ARAF^282-579^ (ARAF) or StrepII-CRAF^324-618^ (CRAF), respectively, in insect cells. The ARAF complex was purified by Ni-NTA followed by gel filtration with a Superdex 200 10/300. The CRAF complex was purified by Ni-NTA, the bound and eluted from StreptactinMacroprep beads, then finished by gel filtration over a Superdex 200 10/300. ARAF^SSDD^ and CRAF^SSDD^ were both prepared by co-expressing MEK1^SASA^ with either his_6_-MBP-TEV-ARAF^274-606, G300S, Y301/302D^ (ARAF^SSDD^) or his6-MBP-TEV-CRAF^314-618, Y340/341D^ (CRAF^SSDD^). In both cases, samples were purified by Ni-NTA, cleaved with TEV protease overnight to remove the MBP tag and re-applied to Ni-NTA beads. This flowthrough was concentrated to 1 ml and injected onto a Superdex 200 10/300 gel filtration column.

Full-length BRAF was co-expressed with MEK1^SASA^ in SF9 insect cells and co-purified with endogenous 14-3-3 proteins. Protein expression and purification proceeded as was standard for the dimeric complexes, harvesting cells 72 hours post infection, and lysing via sonication in Ni-binding buffer (pH 8.0, 50 mM Tris, 150 mM NaCl, 5 mM MgCl_2_, 1 mM TCEP, and 1 μM AMP-PNP). The lysate was ultracentrifuged at 40000 rpm for 1 hour. This lysate was purified by application and elution from Ni-NTA beads and a 5 ml StrepTrap column, eluting with binding buffer supplemented with 300 mM imidazole and 5 mM desthiobiotin, respectively. Purification finished with size exclusion chromatography on a Superose 6 or a Superdex 200 Increase column. Fractions were analyzed by SDS-page, and appropriate fractions were pooled and used for assays.

#### pMEK TR-FRET kinase assays

Inhibition assays were performed using the TR-FRET setup established previously (25, 26). MEK phosphorylation is detected by phospho-MEK antibody binding, and leads to an increase in FRET signal. Inhibitors were again dispensed by HP300e into black 384-well round bottom plates and normalized to 1% DMSO. MEK-B and detection reagent concentrations were maintained at the levels used previously (250 nM MEK-B, 0.5 nM antibody, and 62.5 nM XL665) but due to the lower activities of the monomeric RAF complexes, assays were done using 25 nM enzyme concentration, unless otherwise specified for a particular experiment. Additionally, ATP concentration was kept at a constant 200 μM, again unless otherwise specified for a specific experiment. Reactions were incubated with inhibitor for 40 minutes, allowed to react upon ATP addition for 30 minutes, then incubated with detection reagents for 40 minutes before reading out FRET signal ratio using a PHERAstar microplate reader.

“Standard” dose responses were fit in GraphPad Prism with three or four parameter curves. Inhibitors that produce paradoxical activation were also processed in GraphPad Prism, using the Bell-Shaped Curve Fit to determine IC_50_, AC_50_, and Hill slope values for each. Bell-shaped curves are somewhat more challenging for Prism to fit well, if this is attempted using the default fit settings. We found the most consistent way to produce reasonable looking fits, without applying unnecessarily stringent constraints, is to manually assign roughly accurate initial values for the AC_50_, IC_50_, both plateaus (no inhibitor and fully inhibited FRET signal), and the “dip” (peak FRET achieved by activation). Assays were performed in triplicate three independent times, unless specifically mentioned otherwise.

#### pERK TR-FRET kinase assays

Inhibition assays were conducted using a modified cell-based HTRF phospho-ERK kinase assay kit (Revivty). ERK phosphorylation is detected by concurrent binding of a D2 ERK antibody and a Europium conjugated phospho-ERK antibody, and leads to an increase in FRET signal (665/620). Inhibitors were again dispensed by HP300e into black 384-well round bottom plates and normalized to 1% DMSO. Assays were performed using 6.25 nM BRAF monomer, 250 nM MEK, 500 nM ERK, and 200 μM ATP. Reaction solutions, containing inhibitor, BRAF, MEK, and ERK, were incubated with inhibitor for 40 minutes, allowed to react upon ATP addition for 30 minutes, then incubated with detection reagents at an 80:1 detection buffer to antibody ratio overnight, before measuring FRET signal ratio using a PHERAstar microplate reader.

#### Mass photometry

The BRAF used for mass photometry (MP) measurements was expressed and purified as described in previously (25). Before collecting mass photometry videos, BRAF aliquots were thawed from -80 and centrifuged at high speed (13200 rpm). MP buffer (25 mM HEPES pH 7.0, 2 mM MgCl_2_, 100 mM NaCl, 10 uM ATPγS) was filtered using a 0.1 µm filter before usage. Initial concentration of BRAF was determined by Bradford assay.

Mass photometry measurements were taken of the BRAF:MEK1^SASA^ complex using the Refeyn TwoMP mass photometer with an Accurion vibration-isolation bench. For data acquisition, pre-cleaned Refeyn Sample Carrier Slides were assembled with reusable six-well silicone gasket cassettes, allowing for collection of six measurements before switching to a new coverslip and fresh gasket. Before taking a reading, the well formed by the gasket and coverslip was filled with 16 µl of room temperature MP buffer (25 mM HEPES, 2 mM MgCl_2_, 100 mM NaCl, 10 uM ATPγS), to allow for microscope focusing on the glass. Once focused, with no evidence of air bubbles or debris in the field of view, 2 µl of 580 nM BRAF:MEK1^SASA^ were added to the 16 µl of buffer in the well, and this 18 µl was pipetted up and down four times with a P20 pipette to ensure mixing of macromolecules within the droplet. Measurements were started immediately and recorded for 60 seconds. For measurements taken in presence of inhibitor, all reactions were incubated with the given concentration of inhibitor for 40 minutes at room temperature, with DMSO concentrations manually normalized to 1% for each reaction, regardless of inhibitor concentration. These reactions were set up manually in standard Eppendorf tubes, incubation in 396 well plates of several varieties was not compatible with collection of MP measurements, resulting in erratic readings with high proportions of unbinding events. Contrast values of experimentally observed counts were converted to kDa molecular weights via mass calibrations. Two mass calibration mixes were used in this capacity, providing consistent results with each other: 3 µM Thyroglobulin and 10 µM BSA, or 3 µM Thyroglobulin and 10 µM β-amylase (Sigma β-amylase from sweet potato (A8781), Sigma Bovine Thyroglobulin (609310), and Thermofisher Bovine Serum Albumin Standard (23209)).

In the resulting molecular weight versus counts histograms, a maximum of four discrete peaks were observed (45 kDa, 80 kDa, 115 kDa, 160 kDa), made up of the following complexes: MEK/BRAF/BRAF/MEK (160 kDa), BRAF/BRAF/MEK (115 kDa), BRAF/MEK (80 kDa), BRAF/BRAF (70 kDa), MEK (45 kDa), and BRAF (35 kDa). The counts within each peak were obtained by binning all counts within 15 kDa above and below each peak center (45 ± 15, 80 ± 15, 115 ± 15, 160 ± 15). The 80, 115, and 160 kDa counts were converted into numbers of specific protein molecules based on the stoichiometry of the BRAF:MEK complexes within each population, using equations 1 – 6. For classification of the 80 kDa counts, we assumed all counts are BRAF:MEK heterodimers, due to RAF’s increased affinity for MEK compared to dimerization (27, 39, 40), as indicated in equations 1 and 4. For the 45 kDa bin, the number of counts corresponding to free BRAF protomers versus free MEK protomers was determined by assuming a given total BRAF:MEK stoichiometry in the sample, and calculating the numbers of each that must be in the 45 kDa population to meet the assumed stoichiometry, as shown in equations 7 – 9 and 10a, 11a (1:1 stoichiometry) or 10b,11b (1:1.5 stoichiometry). Once the number of species corresponding to each count was calculated, the percents of 115 kDa BRAF, 160 kDa BRAF and dimeric BRAF were calculated using equations 12 – 17, while the K_d_ values of each binding event were calculated with equations 18 - 21. The BRAF:MEK stoichiometry ratios (1:1 or 1:1.5) used for converting 45 kDa counts to numbers of BRAF and MEK protomers were informed by SDS-PAGE analysis of the BRAF:MEK sample used for MP experiments. The molar concentration of the sample (*conc.BRAF*) used in equations 18 – 21 was determined by Bradford assay. In the text, all percents and K_d_ values referred to were calculated based on a 1:1 BRAF:MEK stoichiometry.

Mass Photometry Equations:

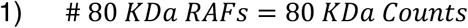

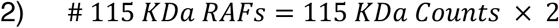

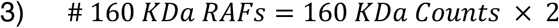

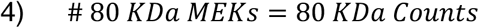

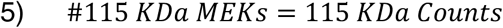

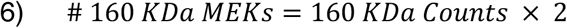

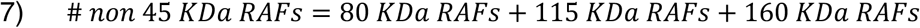

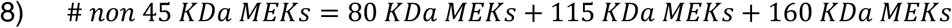

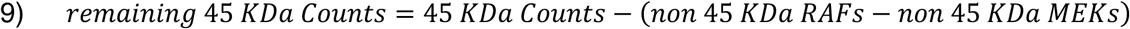

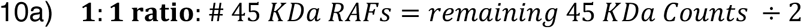

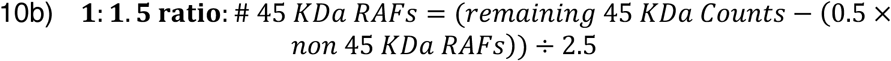

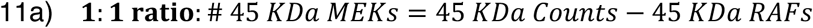

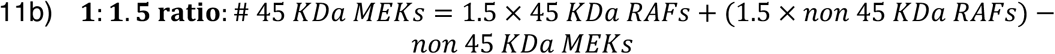

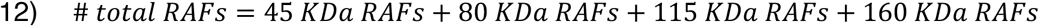

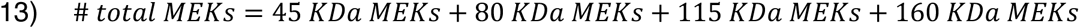

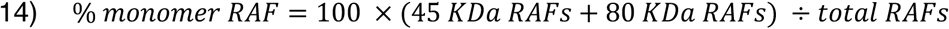

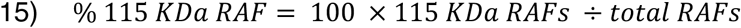

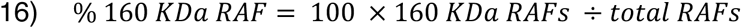

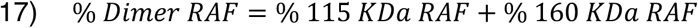

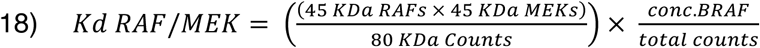

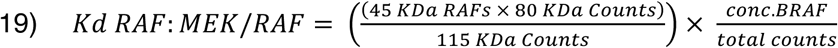

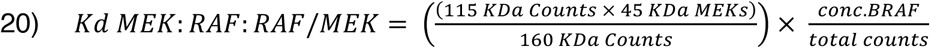

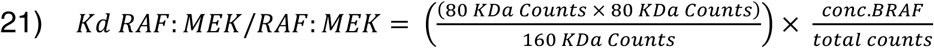

## Data Availability

All data are included in the manuscript and supplemental materials.

## Supporting Information

This article contains supporting information.

## Author Contributions

E. Tkacik, B.H. Ha, J. Vinals, D. Jang and K. Boxer purified proteins for inhibitor characterization. E. Tkacik performed enzyme inhibition assays. M.J. Eck supervised the research. E. Tkacik and M.J. Eck wrote the manuscript. All authors reviewed and approved this manuscript prior to publication.

## Funding and Additional Information

This work was supported in part by NIH grants R35CA242461 (M.J.E.), P50CA165962 (M.J.E.), and by the Dana-Farber Pediatric Low-Grade Astrocytoma Program. E.T. is the recipient of a graduate research fellowship from the Chleck Family Foundation.

