## Supplemental Materials for "In vitro reconstitutions suggest a general model for paradoxical activation of ARAF, BRAF, and CRAF by diverse RAF inhibitor types that does not rely on negative allostery"

Emre Tkacik et al.

This file includes:

Supplemental Tables 1 to 3

Supplemental Figures 1 to 4

**Supplemental Table 1. Effect of ATP concentration on BRAF paradoxical activation parameters.***

| | $AC_{50}$ (nM) | | |
|---|---|---|---|
| **inhibitor** | **10 μM ATP** | **100 μM ATP** | **1000 μM ATP** |
| GDC0879 | 3.31 ± 1.27 | 0.56 ± 0.19 | 6.13 ± 5.97 |
| SB590885 | 2.76 | 0.33 | 2.97 ± 1.97 |
| Tovorafenib | 0.81 ± 0.45 | 0.29 ± 0.33 | 0.44 ± 0.48 |
| Ponatinib | 2.25 ± 1.25 | 0.61 ± 0.61 | 1.29 ± 0.80 |
| AZ628 | 0.44 ± 0.19 | 0.22 ± 0.20 | 0.28 ± 0.14 |
| Belvarafenib | 0.43 ± 0.38 | 0.30 ± 0.07 | 0.69 ± 0.58 |
| LY3009120 | 0.73 ± 0.17 | 0.49 ± 0.17 | 0.78 ± 0.13 |
| | **$IC_{50}$ (nM)** | | |
| **inhibitor** | **10 μM ATP** | **100 μM ATP** | **1000 μM ATP** |
| GDC0879 | 583.7 ± 171.2 | 6051 ± 350.0 | >10000 |
| SB590885 | 76.20 ± 87.55 | 564.9 ±112.9 | 6423 ± 506.3 |
| Tovorafenib | 257.3 ± 27.95 | 206.3 ± 10.11 | 167.2 ± 53.43 |
| Ponatinib | 119.9 ± 57.93 | 103.3 ± 2.47 | 97.76 ± 46.62 |
| AZ628 | 28.83 ± 5.61 | 24.39 ± 1.40 | 31.60 ± 5.02 |
| Belvarafenib | 36.64 ± 17.78 | 26.75 ± 2.97 | 37.01 ± 10.48 |
| LY3009120 | 36.58 ± 0.42 | 39.81 ± 4.19 | 39.80 ± 2.68 |

* Activation ($AC_{50}$) and inhibitory ($IC_{50}$) fitting parameters are reported as mean ± SD from two or more independent experiments performed in triplicate (n≥3). Representative concentration response curves for a subset of inhibitors are presented in Figure 4 of the main text. Standard error for SB590885 is not always reported because for some of the independent experiments, $AC_{50}$ could not be determined.

**Supplemental Table 2. Effect of RAF inhibitor concentrations on BRAF/MEK dissociation constants as determined by mass photometry.***

| BRAF/MEK Stoichiometry (1:1) | | | | |
|---|---|---|---|---|
| **GDC0879** | | | | |
| **[inhibitor] (nM)** | **$K_{d, R\text{-}M}$ (nM)** | **$K_{d, RM\text{-}R}$ (nM)** | **$K_{d, MRR\text{-}M}$ (nM)** | **$K_{d, RM\text{-}RM}$ (nM)** |
| 100000 | 57 ± 23 | 11 ± 4 | 31 ± 6 | 6 ± 2 |
| 10000 | 55 ± 15 | 21 ± 6 | 22 ± 3 | 9 ± 5 |
| 1000 | 53 ± 43 | 27 ± 13 | 24 ± 3 | 14 ± 5 |
| 100 | 27 ± 26 | 150 ± 64 | 44 ± 10 | 358 ± 170 |
| 10 | 16 ± 15 | 124 ± 82 | 46 ± 9 | 448 ± 163 |
| 1 | 33 ± 26 | 137 ± 57 | 48 ± 13 | 327 ± 256 |
| 0.1 | 18 ± 7 | 163 ± 54 | 51 ± 20 | 480 ± 193 |
| 0.01 | 19 ± 7 | 205 ± 84 | 45 ± 18 | 461 ± 179 |
| **Vemurafenib** | | | | |
| **[inhibitor] (nM)** | **$K_{d, R\text{-}M}$ (nM)** | **$K_{d, RM\text{-}R}$ (nM)** | **$K_{d, MRR\text{-}M}$ (nM)** | **$K_{d, RM\text{-}RM}$ (nM)** |
| 100000 | 20 ± 18 | 164 ± 89 | 253 ± 189 | 2089 ± 760 |
| 10000 | 13 ± 8 | 169 ± 76 | 179 ± 18 | 2509 ± 678 |
| 1000 | 27 ± 27 | 227 ± 143 | 144 ± 28 | 1758 ± 977 |
| 100 | 22 ± 12 | 178 ± 89 | 77 ± 29 | 586 ± 161 |
| 10 | 20 ± 13 | 116 ± 72 | 59 ± 28 | 350 ± 100 |
| 1 | 16 ± 11 | 182 ± 107 | 36 ± 8 | 504 ± 250 |
| 0.1 | 37 ± 38 | 172 ± 129 | 39 ± 7 | 458 ± 437 |
| 0.01 | 56 ± 58 | 223 ± 36 | 47 ± 22 | 259 ± 110 |
| **Naporafenib** | | | | |
| **[inhibitor] (nM)** | **$K_{d, R\text{-}M}$ (nM)** | **$K_{d, RM\text{-}R}$ (nM)** | **$K_{d, MRR\text{-}M}$ (nM)** | **$K_{d, RM\text{-}RM}$ (nM)** |
| 100000 | 69 ± 31 | 20 ± 8 | 70 ± 23 | 22 ± 8 |
| 10000 | 61 ± 38 | 19 ± 11 | 58 ± 37 | 20 ± 14 |
| 1000 | 40 ± 34 | 25 ± 27 | 49 ± 28 | 45 ± 67 |
| 100 | 34 ± 26 | 115 ± 65 | 64 ± 43 | 231 ± 101 |
| 10 | 45 ± 60 | 143 ± 77 | 62 ± 39 | 328 ± 169 |
| 1 | 30 ± 24 | 114 ± 37 | 81 ± 42 | 375 ± 180 |
| 0.1 | 21 ± 16 | 107 ± 36 | 67 ± 18 | 421 ± 214 |
| 0.01 | 37 ± 49 | 179 ± 157 | 54 ± 10 | 365 ± 167 |

*$K_d$ values for the indicated reactions (nM) are reported as mean ± SD from three independent measurements at each inhibitor concentration. Dissociation constants correspond to the scheme in Figure 5B in the main text, where R represents BRAF and M represents MEK1, and the "-" indicates the association-dissociation reaction. Corresponding calculated species abundances are provided in Supplemental Table 3.

**Supplemental Table 3. Effect of RAF inhibitor concentrations on percent abundance of BRAF/MEK1 species as determined by mass photometry.***

| BRAF/MEK Stoichiometry (1:1) | | | | |
|---|---|---|---|---|
| **GDC0879** | | | | |
| **[inhibitor] (nM)** | **% Monomer** | **% 115 kDa** | **% 160 kDa** | **% Dimer** |
| 100000 | 51 ± 11 | 31 ± 6 | 18 ± 5 | 49 ± 11 |
| 10000 | 62 ± 4 | 22 ± 4 | 17 ± 1 | 38 ± 4 |
| 1000 | 65 ± 15 | 21 ± 10 | 15 ± 5 | 35 ± 15 |
| 100 | 88 ± 6 | 9 ± 5 | 3 ± 1 | 12 ± 6 |
| 10 | 84 ± 7 | 12 ± 6 | 4 ± 1 | 16 ± 7 |
| 1 | 88 ± 6 | 9 ± 5 | 3 ± 1 | 12 ± 6 |
| 0.1 | 89 ± 2 | 8 ± 3 | 3 ± 1 | 11 ± 2 |
| 0.01 | 90 ± 4 | 7 ± 3 | 3 ± 1 | 10 ± 4 |
| **Vemurafenib** | | | | |
| **[inhibitor] (nM)** | **% Monomer** | **% 115 kDa** | **% 160 kDa** | **% Dimer** |
| 100000 | 90 ± 5 | 9 ± 5 | 1 ± 1 | 10 ± 5 |
| 10000 | 90 ± 3 | 9 ± 3 | 1 ± 0 | 10 ± 3 |
| 1000 | 92 ± 4 | 7 ± 4 | 1 ± 1 | 8 ± 4 |
| 100 | 90 ± 7 | 8 ± 7 | 2 ± 1 | 10 ± 7 |
| 10 | 83 ± 12 | 13 ± 8 | 4 ± 3 | 17 ± 12 |
| 1 | 89 ± 5 | 8 ± 4 | 3 ± 0 | 11 ± 5 |
| 0.1 | 88 ± 11 | 10 ± 10 | 3 ± 1 | 12 ± 11 |
| 0.01 | 93 ± 4 | 5 ± 2 | 2 ± 1 | 7 ± 4 |
| **Naporafenib** | | | | |
| **[inhibitor] (nM)** | **% Monomer** | **% 115 kDa** | **% 160 kDa** | **% Dimer** |
| 100000 | 66 ± 11 | 26 ± 7 | 8 ± 4 | 34 ± 11 |
| 10000 | 60 ± 15 | 28 ± 11 | 12 ± 5 | 40 ± 15 |
| 1000 | 56 ± 21 | 30 ± 13 | 14 ± 9 | 44 ± 21 |
| 100 | 84 ± 9 | 12 ± 6 | 4 ± 3 | 16 ± 9 |
| 10 | 87 ± 7 | 10 ± 6 | 3 ± 2 | 13 ± 7 |
| 1 | 87 ± 5 | 11 ± 4 | 3 ± 2 | 13 ± 5 |
| 0.1 | 85 ± 5 | 12 ± 3 | 3 ± 1 | 15 ± 5 |
| 0.01 | 87 ± 8 | 10 ± 6 | 3 ± 2 | 13 ± 8 |
| 0 | 79 ± 11 | 15 ± 8 | 5 ± 3 | 21 ± 11 |

*Fractional species abundances are reported as mean percent ± SD from three independent measurements at each inhibitor concentration. Calculated abundance of each molecular-weight species was used to calculate dissociation constants shown in Supplemental Table 2.

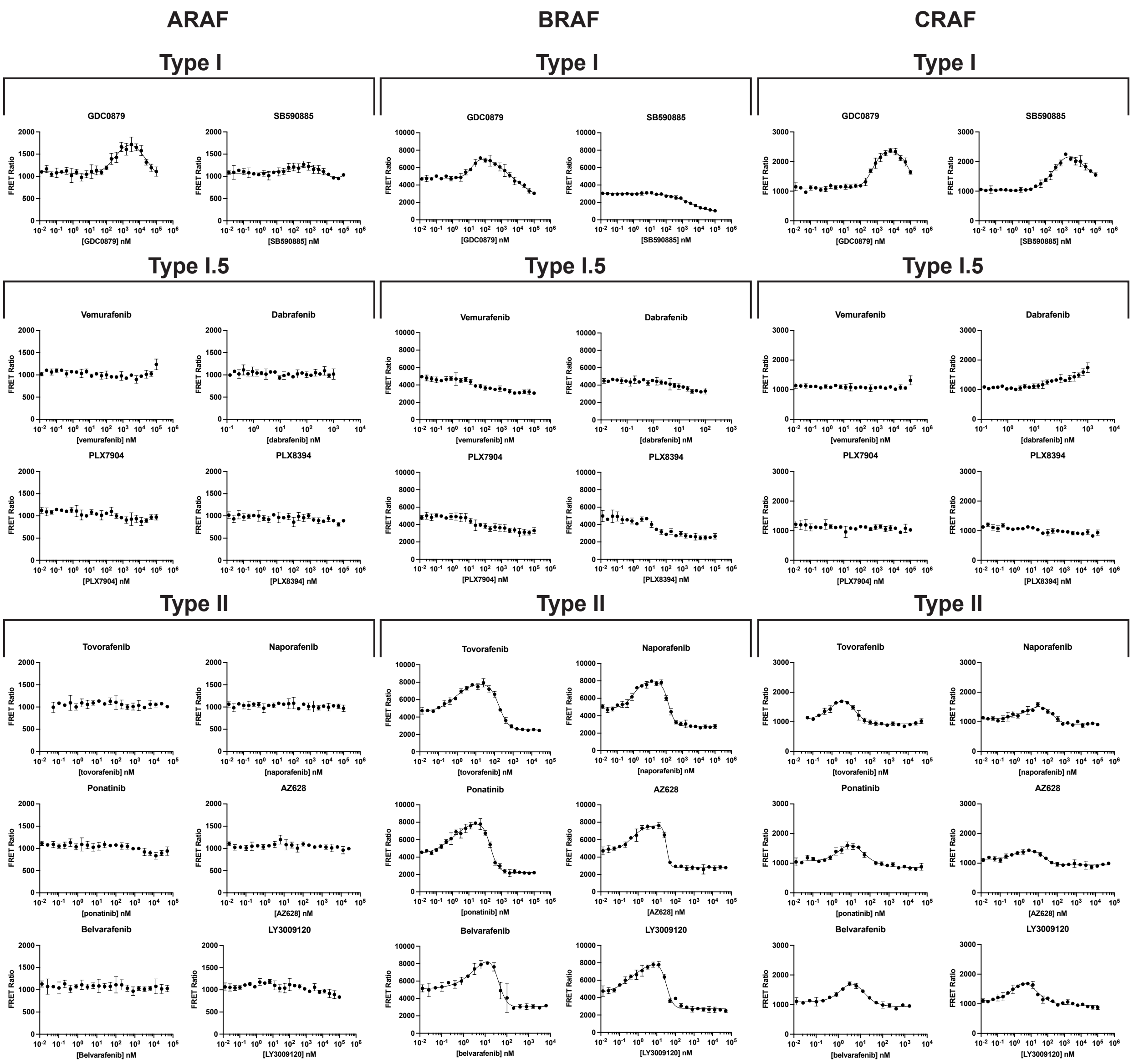


**Supplemental Figure 1. Concentration-response curves for RAF inhibitors with ARAF, BRAF, and CRAF monomer complexes.** Representative concentration-response curves for the inhibitors indicated are shown for ARAF, BRAF, and CRAF monomer complexes, with data plotted as mean ± SD from one independent experiment performed in triplicate (n = 3). Inhibitors are grouped by class according to binding mode, with ARAF dose response curves on the left, BRAF dose response in the middle, and CRAF on the right.

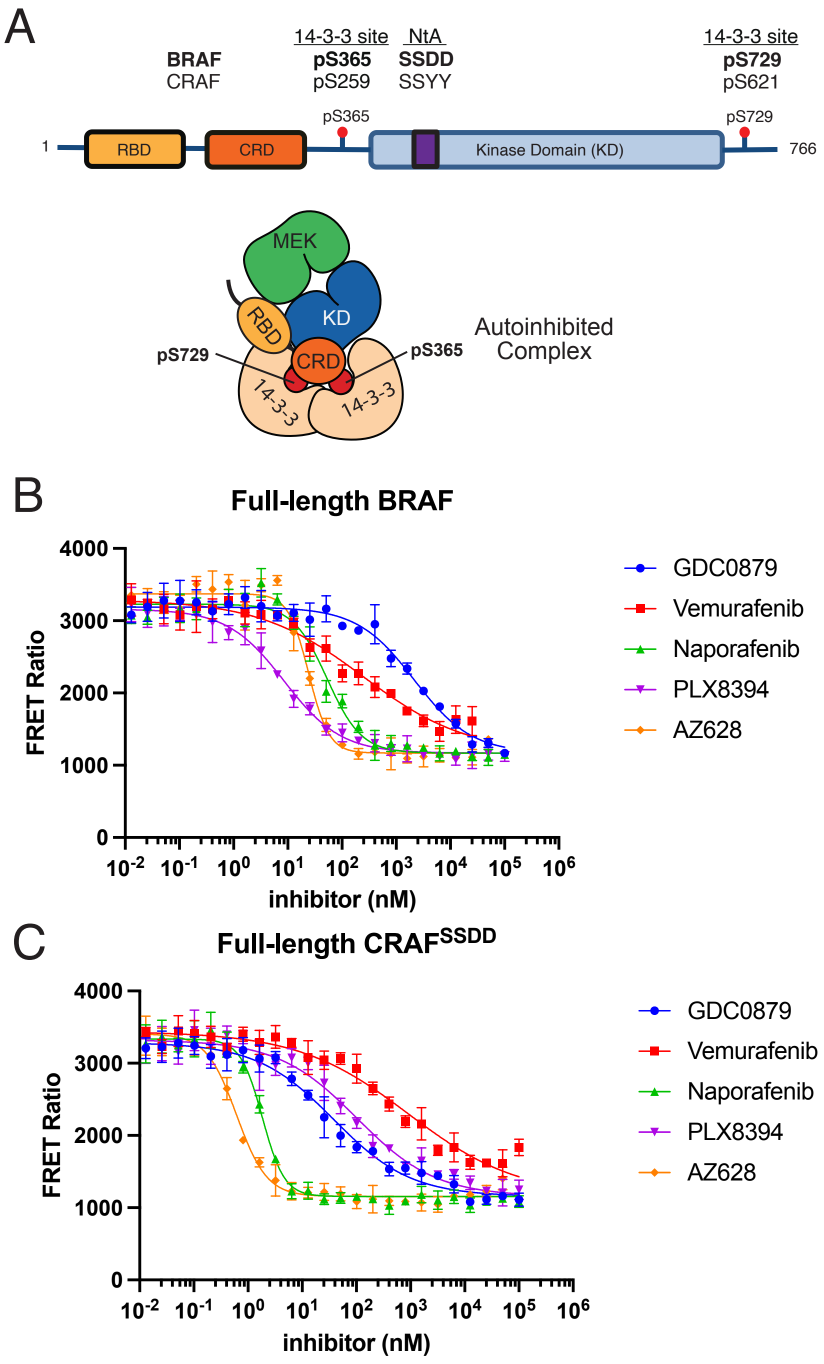


**Supplemental Figure 2. Concentration response curves for titration of full-length autoinhibited BRAF/MEK1/14-3-3 and CRAF/MEK1/14-3-3 complexes with RAF inhibitors.** A, “Linear” RAF domain schematic and schematic depicting the three-dimensional organization of autoinhibited BRAF and CRAF. B and C, Concentration-response curves for full-length complexes of BRAF/MEK1/14-3-3 (B) or CRAF/MEK1/14-3-3 (C) titrated with the indicated RAF inhibitors. The CRAF complex was prepared with CRAF$^{SSDD}$. The low activity present in these autoinhibited preparations may stem, at least in part, from minor contamination with active dimers. Note the lack of paradoxical activation. Data are plotted as mean ± SD from three independent experiments performed in triplicate (n=3).

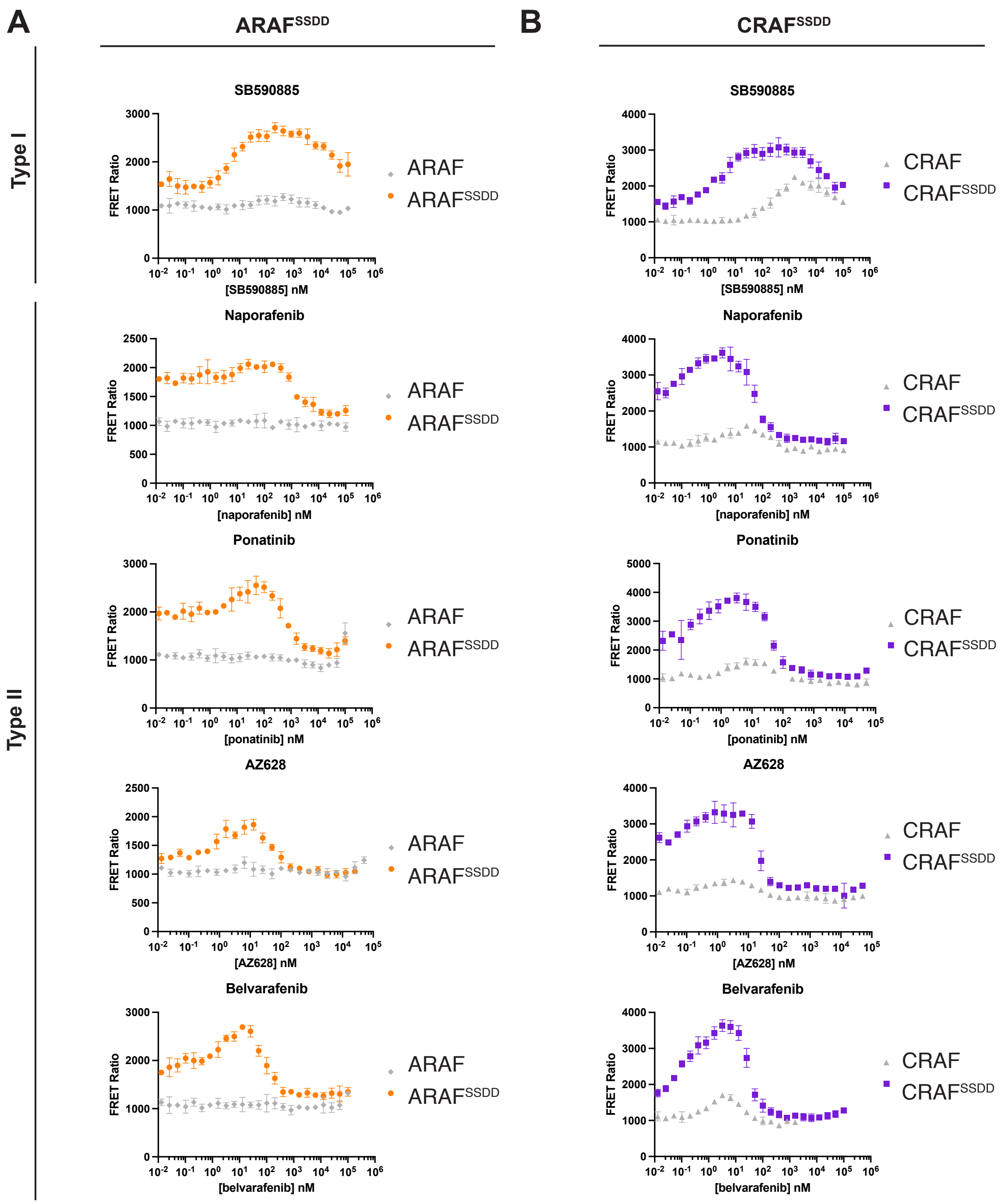


**Supplemental Figure 3.** A, Concentration-response curves for titration of ARAF$^{SSDD}$/MEK1 (ARAF$^{SSDD}$) and CRAF$^{SSDD}$/MEK1 (CRAF$^{SSDD}$) with the indicated RAF inhibitors. B, Concentration-response curves for titration of CRAF$^{SSDD}$ with the indicated RAF inhibitors. Data for the wild-type ARAF and CRAF are reproduced from Supplemental Figure 1 to facilitate comparison and are plotted in gray. Curves for additional type I and type II inhibitors are shown in Figure 3 in the main text. Representative experiments are plotted as mean ± SD from one independent experiment performed in triplicate (n=3).

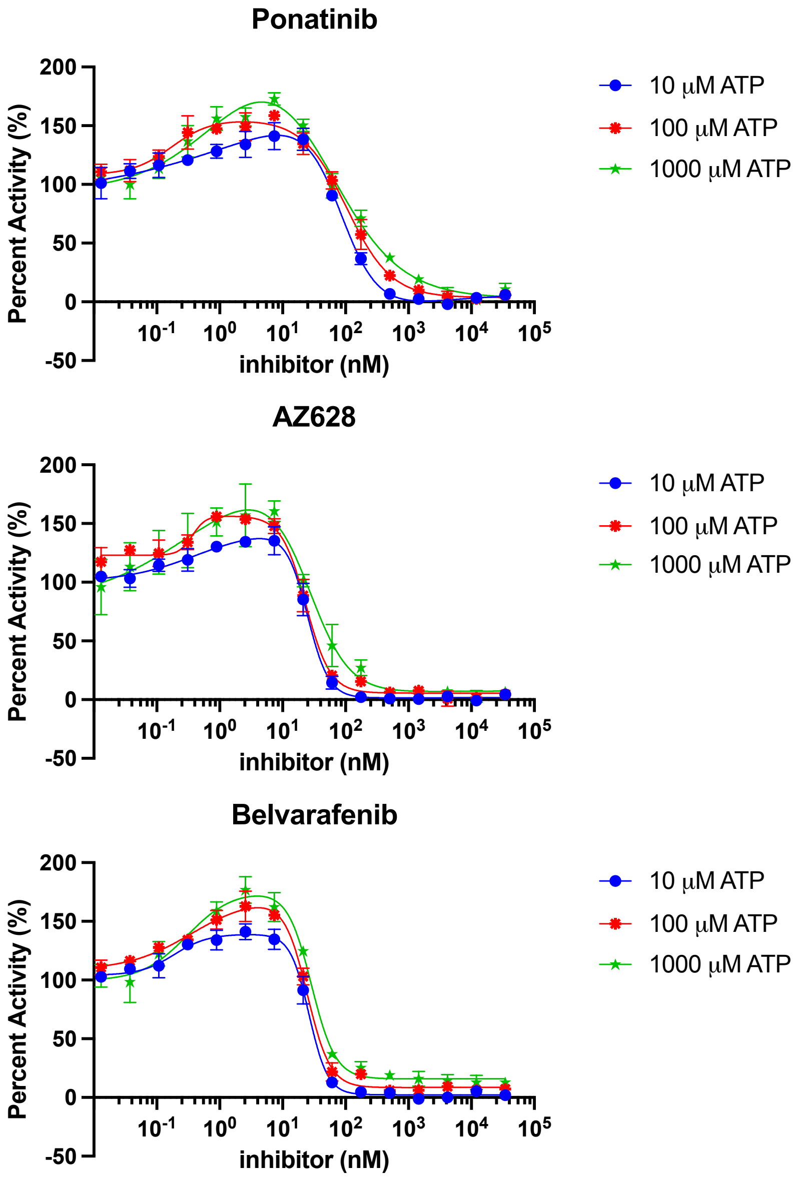


**Supplemental Figure 4. Effect of ATP concentration on paradoxical activation of BRAF by additional type II inhibitors.** Concentration-response curves measuring MEK phosphorylation upon treatment of the BRAF monomer complex with increasing concentrations of the indicated inhibitor at ATP concentrations of 10 µM (blue), 100 µM (red), and 1000 µM (green). MEK phosphorylation activity is plotted as percent activity, normalized to the activity in the absence of inhibitor at each ATP concentration. Data points are plotted as mean ± SD from one independent experiment in triplicate (n≥3).
